# Diel transcriptional pattern contributes to functional and taxonomic diversity in supraglacial microbial communities

**DOI:** 10.1101/2021.01.18.427117

**Authors:** Francesca Pittino, Simone Zordan, Roberto S. Azzoni, Guglielmina Diolaiuti, Roberto Ambrosini, Andrea Franzetti

## Abstract

Despite the harsh environmental conditions, glacier surfaces host metabolically active bacterial communities, especially in cryoconite holes, small ponds filled with melting water and with a fine-grained sediment at the bottom. We investigated the daily changes in transcript profiles of the microbial community of a cryoconite hole on an Alpine glacier. Using a metatranscriptomic shotgun sequencing, we observed different level of expression of the main carbon and energy metabolisms along the day. Oxygenic and anoxygenic photosynthesis peaked their activity at the sunrise and sunset, respectively, and showed an inhibition at midday, in response to high solar radiation. Carbon fixation genes were expressed all day long with the lowest coverage at night. Different microbial populations were responsible for this metabolic function along the day. Cyanobacteria and Algae were the most active primary producers at the sunrise and the sunset, whereas at night and at noon chemosynthetic proteobacteria, likely hydrogen oxidisers, were most active. Furthermore, the observed temporal cascade of transcript peaks of photosynthesis and respiration recalls those occurring in both coastal and open waters in ocean, thus supporting the hypothesis that conserved temporally phased biotic interactions are ubiquitous among aquatic communities worldwide.

## INTRODUCTION

Cryoconite holes are small ponds filled with of meltwater with a sediment at the bottom present on the surface of most glaciers. Although they are characterized by extreme conditions such as low temperatures and high solar irradiance, they host bacterial communities with high taxonomic and functional biodiversity (Cook *et al.*, 2015). It has been supposed that such high functional diversity could be due to the high versatility of some of the most abundant populations (Darcy *et al.*, 2011). In a previous investigation, we demonstrated that these supraglacial communities exhibit high functional biodiversity since they exploit organic matter both as energy and carbon source, and use both oxygenic and anoxygenic photosynthesis with pure autotrophic and mixotrophic lifestyles (Franzetti *et al.*, 2016). Due to the lack of expression studies in these environments, it is currently unknown whether a diel temporal pattern of expression occurs and how it may contribute to the overall functional and taxonomic biodiversity. To fill this gap of knowledge, we collected four cryoconite samples from a single hole along a clean summer day (4.30 am, 7.30 am, 1.30 pm and 7.30 pm) on the Forni Glacier (Italy) on 24 July 2018 and used shotgun metatransctriptomics sequencing to investigate the expression of the main metabolic functions along the day. We focused on carbon and energy metabolisms by comparing the total coverage of marker gene transcripts for photosynthesis, use of inorganic and organic compounds as energy source, respiration and autotrophy/heterotrophy.

## RESULTS

Table 1 reports the marker gene transcripts whose coverage (mean number per base of reads mapping the genes) was used to infer the expression of each metabolism and their normalized coverages.

**Table 1.** Transcript coverages of the marker genes.

| Metabolism | Gene | Short name | KO Orthology | Coverage |  |  |  |
| --- | --- | --- | --- | --- | --- | --- | --- |
|  |  |  |  | 4.30 am | 7.30 am | 1.30 pm | 7.30 pm |
| **Oxygenic photosynthesis** | photosystem II P680 reaction center D2 protein | *psbD* | K02706 | 2.1 | 317.4 | 83.8 | 163.5 |
| **Aerobic anoxygenic photosynthesis** | photosynthetic reaction center L and M subunits | *pufM* | K08929 | 3.2 | 32.4 | 0.0 | 69.6 |
| **CO<sub>2</sub> fixation, Calvin-Benson cycle** | ribulose bisphosphate carboxylase small and large chain (RubisCO) | *rbcL* | K01602, K01601 | 25.5 | 96.8 | 84.2 | 50.0 |
| **Chitin assimilation** | chitinase | *chiA* | K01183 | 68.7 | 6.5 | 0.0 | 35.1 |
| **Hydrolysis O- and S-glycosyl compounds** | endo-1,4-beta-xylanase | *xynA* | K01181 | N.D | N.D | N.D | N.D |
| **Carbon monoxide oxidation** | carbon-monoxide dehydrogenase medium and large subunits | *coxML* | K03519, K03520 | N.D | N.D | N.D | N.D |
| **Aerobic respiration** | cytochrome c oxidase subunit I | *coxA* | K02274 | 908.8 | 283.4 | 677.6 | 735.1 |
| **Dissimilatory nitrate reduction (Denitrification/Nitrification)** | nitrate reductase alpha subunit | *narG* | K00370 | 748.0 | 145.8 | 568.0 | 454.8 |
| **Dissimilatory nitrate reduction (Denitrification)** | nitrite reductase (NO-forming) [EC:1.7.2.1] | *nirK* | K00368 | 831.3 | 129.3 | 778.2 | 408.5 |
| **Dissimilatory nitrate reduction (Denitrification)** | nitric oxide reductase subunit B [EC:1.7.2.5] | *norB* | K04561 | 815.5 | 105.8 | 666.3 | 409.1 |
| **Ammonia oxidation (nitrification)** | ammonia monooxygenase A and B subunits | *amoA* | K10944 | 0.0 | 1.0 | 0.1 | 0.1 |
| **Sulfur oxidation** | sulfur-oxidizing protein | *sox A* | K17222 | 0.0 | 13.3 | 0.0 | 40.3 |
| **Hydrogen oxidation** | hydrogenase large subunit | *hya B* | K06281 | 19.0 | 0.0 | 26.0 | 12.5 |
| **Nitrogen fixation** | nitrogenase iron protein | *nifH* | K02588 | 0.2 | 0.0 | 0.3 | 2.4 |

We also used transcript sequences for the taxonomic attribution of microorganisms expressing specific metabolic genes (SM1). Finally, we report the transcript coverage of all the annotated KO (KEGG Orthology) molecular functions (SM2).

## DISCUSSION

Results showed that carbon and energy metabolisms had different patterns of expression along the day (Figure 1, Table 1). Before sunrise (4.30 am) aerobic respiration and denitrification were the dominant energy metabolisms, while neither oxygenic nor anoxygenic phototrophy were active (Figure 1).

**Figure 1.**
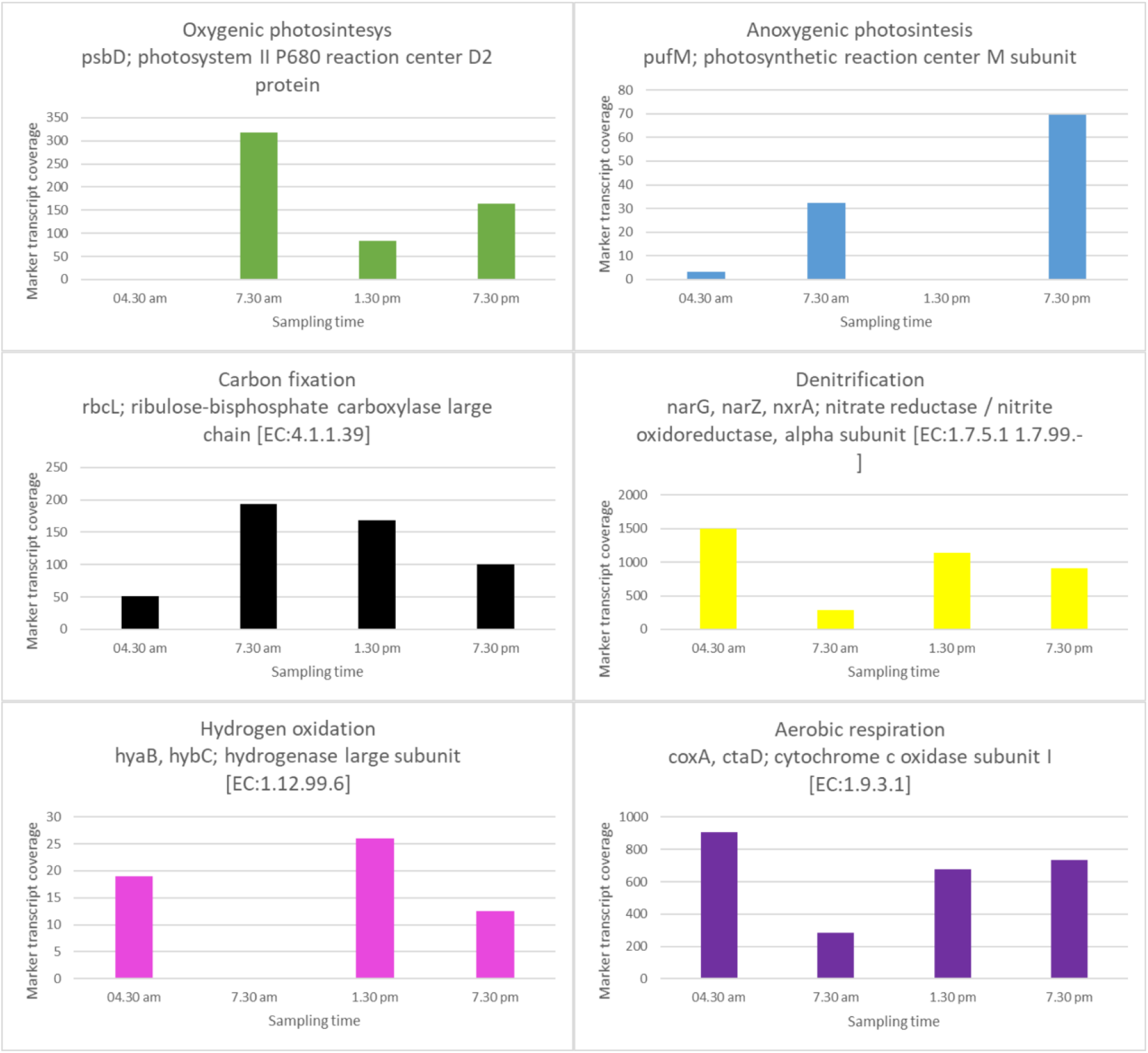
Marker transcript coverages cryoconite. *psb***D**, photosystem II P680 reaction center D2 protein; *puf***M**, photosynthetic reaction center M subunit; *rbc***L**, ribulose bisphosphate carboxylase large chain (RubisCO); *nar***G**; nitrate reductase alpha subunit; *hya***B**: hydrogenase large subunit; *cox***A**: cytochrome c oxidase subunit **I**.

According to the taxonomic affiliation of *cox*A and *nir*K, Actinobacteria and Proteobacteria were the most active taxa involved in respiration with slightly different abundances along the day (SM1). Cyanobacterial respiration was active after sunrise (7.30 am) (20% of abundance) and before the sunset (7.30 pm) (6% of abundance) (Figure 1).

Anaerobic respiration also actively occured in cryoconite holes, seems restricted to denitrification since neither iron reduction (rusticyanin) nor dissimilatory sulfite reductase gene (*dsr*A) were actively transcribed (SM2). Consistently, the development of an anoxic zone 2 mm deep has been recently described in cryoconite (Poniecka *et al.*, 2018) and the complete denitrification pathway was also active (*nar*G, *nir*K, *nor*B), as already reported in the cryoconite of a Chinese glacier (Segawa *et al.*, 2014).

As expected, Cyanobacteria and algae were the active oxygenic phototrophs. Aerobic anoxygenic phototrophs (APPs) resulted affiliated to alpha and beta-proteobacteria (SM1) and are known to be photoheterotrophic, thus using organic matter as carbon source and complementing their energy demand with light (Koblížek, 2015). Both oxygenic (*psb*D) and aerobic anoxygenic (*puf*M) photosynthesis showed reduced activity at the time of the highest solar irradiance (1.30 pm). For oxygenic phototrophs, this temporal pattern is consistent with the described downregulation of PSI and PSII (*psa* and *pdb* genes) of cyanobacteria after the shift from low to high light intensity (Hihara *et al.*, 2001; Ogawa *et al.*, 2018) as a response to light damages of the photosynthetic apparatus. Expression of oxygenic (*psb*D) and aerobic anoxygenic (*puf*M) photosynthesis peaked after sunrise (7.30 am) and before sunset (7.30 pm), respectively (Figure 1). Interestingly, this pattern of the peak of activity of oxygenic photoautotrophs followed by anoxygenic ones and by the oxidative phosphorylation transcript maxima, recalls those occurring in both coastal and open waters in the Pacific ocean (Aylward *et al.*, 2015).

A relevant expression of the CO_2_ fixation genes (*rblc*L) was observed in all the sampling times with the maximum expression after sunrise and the lowest at night. Interestingly, the taxa expressing this metabolic function are extremely variable along the day. Indeed, Cyanobacteria and algae were the most active CO_2_ fixators at 7.30 am and 7.30 pm. Cyanobacteria were not expressing RubisCO gene at 1.30 pm, in disagreement with previous reports showing an up-regulation of this gene under high light intensity (Hihara *et al.*, 2001; Ogawa *et al.*, 2018). At this time of the day, Proteobacteria and algae were the active primary producers. The fact that at night and at 1.30 pm phototrophs were not the most active primary producers, suggests that chemolithoautotrophic microorganisms could be active. Ammonia seems not among the possible inorganic electron donors, because the expression of *amo*A (ammonia monooxygenase A subunit) was negligible at any time, while the oxidation of nitrite to nitrate may be the active chemolitoautrophic pathway because nitrate reductase alpha subunit (*nar*G) was active. However, NarG might catalyse both the oxidation of nitrite and the dissimilatory reduction of nitrate. Moreover, we observed a negligible expression of nitrogenaes (*nif*H), which suggests that nitrogen was not fixed. Therefore, a short nitrogen cycle comprising of dissimilatory nitrate reduction to nitrite and lithotrophic oxidation of nitrite to nitrate could be put forward as a mechanism sustaining the chemosynthesis in these environments. However, the genes for the complete denitrification pathway (*nar*G, *nir*K, *nor*B) showed similar expression levels at all times, suggesting that NarG had reductive catalytic activity rather than oxidative. Carbon monoxide oxidation to CO_2_ was previously hypothesized as a possible lithotrophic metabolism in cryoconite due to the high abundance of carbon-monoxide dehydrogenase (*cox*ML) (Franzetti *et al.*, 2016). However, the transcripts of this gene were not retrieved in this study, thus suggesting, as previously reported (Haan *et al.*, 2001), that carbon monoxide is produced by photochemical reactions in presence of snow. Among the different key genes for lithotrophic sulfur oxidation (*apr*A, *dsr*A), only *sox*A was transcribed (SM2). However, *sox* genes were likely harboured by anoxygenic phototrophs (Dahl, 2008), since they were retrieved only at 7.30 am and 7.30 pm when also the *puf*M genes were highly transcribed and when the primary production was mainly due to phototrophs. Therefore, we excluded sulfur oxidation as an active chemosynthetic process.

Interestingly, hydrogenase (*hya*B) genes were actively transcribed (Figure 1, Table 1) when CO_2_ fixation was due to non-phototrophs (4.30 am and 1.30 pm). This might indicate that hydrogen oxidation might provide the reducing power for primary production during the night and when photosynthesis is inhibited. This hypothesis is supported by the fact that most of *rblc*L transcripts at 4.30 am and 1.30 pm were taxonomically assigned to *Paracoccus* spp. and *Hydrogenophilus* spp., respectively (SM2), which are known for their capacity to use molecular hydrogen as an electron donor (Hayashi *et al.*, 1999; Bardischewsky and Friedrich, 2001). Hydrogen can have both biotic or abiotic origin. Biological hydrogen production could be of both fermentative and photosynthetic origin and is inhibited by oxygen (Benemann, 1997; Das and Veziroǧlu, 2001). Therefore, these processes could be active in the anoxic layer of the cryoconite (Poniecka *et al.*, 2018). Abiotic hydrogen production has been proposed as a mechanism sustaining lithoautotrophic life in deep and subglacial environments. Abiogenic H_2_ is formed due to a variety of water–rock reactions (Telling *et al.*, 2015). Telling and colleagues reported that sufficient H_2_ is produced to support methanogenesis under a Greenland glacier (Telling *et al.*, 2015). Moreover, recent studies demonstrated that cryoconite is among the most radioactive environmental matrices, with activity concentrations exceeding 10,000 Bq kg-1 for single radionuclides (Łokas *et al.*, 2016; Baccolo *et al.*, 2019). This radioactivity might cause radiolytic H_2_ production in the cryoconite holes, which, in turn, supports bacterial metabolism as already reported in deep subsurface environments (Lin *et al.*, 2005). Further studies are required to confirm the presence of hydrogen in cryoconite and to assess its origins.

This first report of shotgun metatranscriptomics in cryoconite therefore revealed a circadian trend of the main energy and carbon metabolisms, which may contribute to the overall functional and taxonomic diversity of these environments (Figure 2).

**Figure 2.**
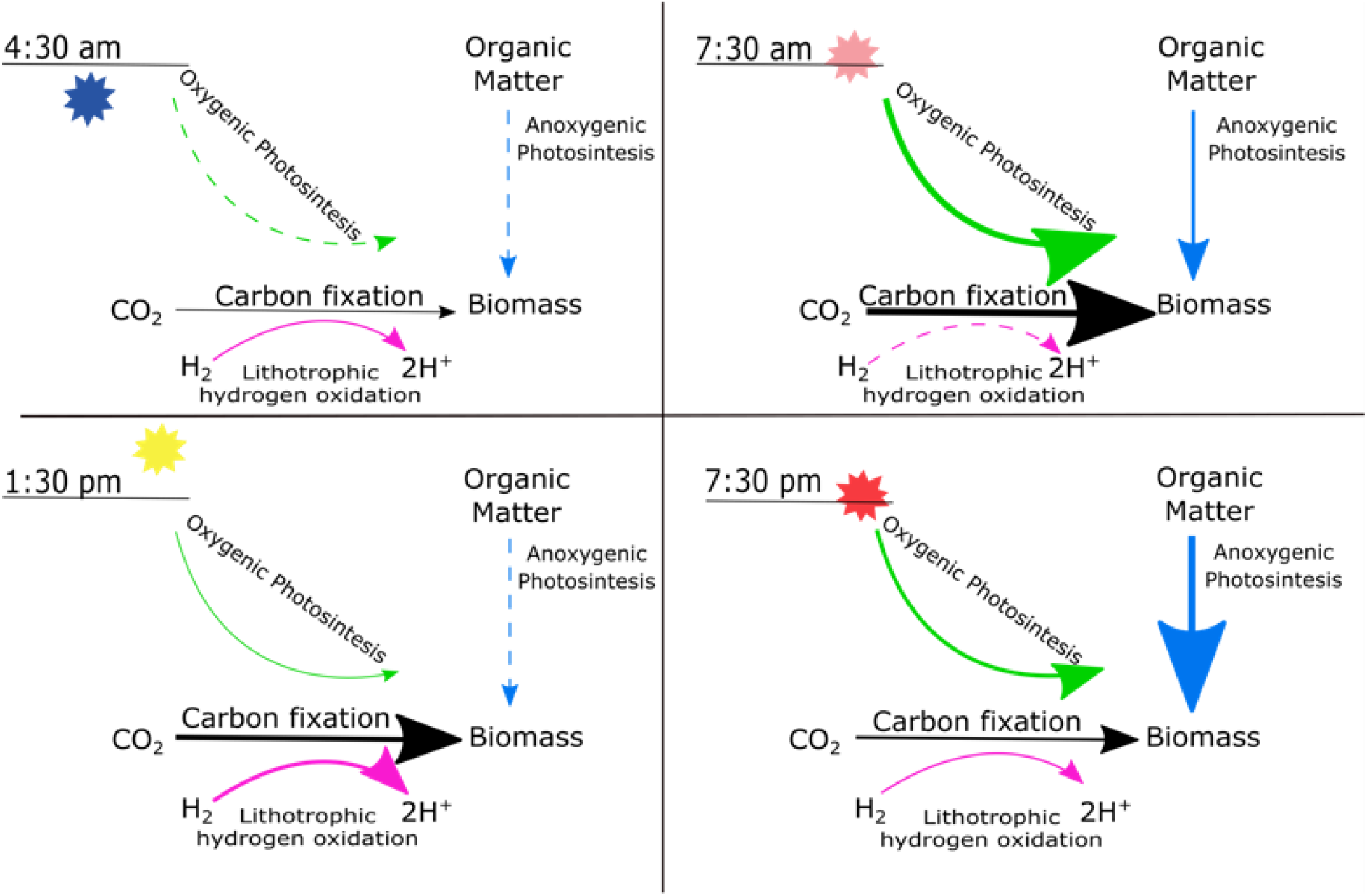
Proposed model of a temporal trend of the main energy and carbon metabolisms. Arrows thickness is proportional to transcript abundances.

Different microbial taxa contribute to the same ecosystem functions (e.g. carbon fixation and respiration) along the day and primary productivity was supported by both light and chemosynthesis. This functional redundancy likely enhances the ecosystem stability in such extreme environments. Results from this study also provide pieces of evidence supporting the hypothesis that conserved microbial community transcriptional and biotic interactions, which follow a daily temporal trend, are ubiquitous among aquatic microbial communities worldwide (Ottesen *et al.*, 2014; Aylward *et al.*, 2015).

## EXPERIMENTAL PROCEDURES

### Sampling and metagenome sequencing

Cryoconite was collected from a single cryoconite hole at four different times (4.30 am, 7.30 am, 1.30 pm, 7.30 pm) on the 24th of July 2018. The cryoconite hole was located on the central ablation tongue of the Forni Glacier (Italy, 46.39868 ° N, 10.58664 ° E) at 2670 m a.s.l. To preserve the RNA content, cryoconite was immediately mixed with a RNA preservative solution (1:5 volume ratio) and stored at −80° within 48 hours. RNA preservative solution was prepared with 4 mL of EDTA 0.5 M, 2.5 mL of Na_3_C_6_H_5_O_7_ 1M, 70 g of (NH_4_)_2_SO_4_, DEPC-H_2_O for a final volume of 100 mL (Gray *et al.*, 2013). Total RNA was extracted from 1 g of cryoconite with the RNeasy PowerSoil Total RNA Kit (QIAGEN, Hilden, Germany) according to the manufacturer instructions. After the extraction, residual DNA was removed with the RQ1 RNase-free DNase (Promega) from 6 μL of extracted RNA. The total DNA-free RNA was then treated with the MICROBExpress™ Bacterial mRNAEnrichment Kit (Ambion) and 8 μL of the depleted RNA were retrotranscribed with the RevertAid First Strand cDNA Synthesis Kit (Thermo Scientific, Waltham, MA, USA) using random primers. Shotgun sequencing was applied to cDNA by HiSeq Illumina (Illumina, Inc., San Diego, CA, USA) using a 150 bp x 2 paired-end protocol on one lane. The paired-end reads were quality-trimmed (minimum length: 80 bp; minimum average quality score: 30) using Sickle (https://github.com/najoshi/sickle). Sequence data were submitted to European Nucleotide Archive (ENA), study accession number PRJEB34670 (http://www.ebi.ac.uk/ena/data/view/ PRJEB34670).

### Bioinformatics procedures

Filtered reads were co-assembled using IDBA-UD (Peng *et al.*, 2012) (Peng *et al.*, 2012). IDBA-UD iterated the value of k_mer_ from 40 to 99 (with a step of 5). Predicted genes were inferred from contigs with Prodigal (Hyatt *et al.*, 2010) (Hyatt *et al.*, 2010). KO numbers were assigned to the predicted proteins using the on line tool GhostKOALA (Kanehisa *et al.*, 2016) (Kanehisa *et al.*, 2016). Lowest Common Ancestor (LCA) algorithm was applied to infer taxonomic affiliation of predicted genes using MEGAN default parameters (Huson *et al.*, 2011) (Huson *et al.*, 2011). Hierarchical taxonomic data were visualised with Krona (Ondov *et al.*, 2011) (Ondov *et al.*, 2011). Average per-base coverage of predicted genes was calculated using filtered reads with bowtie2 (Langmead and Salzberg, 2012) (Langmead and Salzberg, 2012), SAMtools (Li *et al.*, 2009) (Li *et al.*, 2009) and bedtools (Quinlan and Hall, 2010) (Quinlan and Hall, 2010). To normalize the different sequencing depth across the samples, sum of gene coverages was normalized to 600,000 for each sample.

## ACKNOWLEDGEMENTS

Authors thank Parco dello Stelvio (Italy) for logistic assistance. This work was partially funded by the University of Milano-Bicocca (PhD fellowship to FP). Bioinformatics analyses have been run on MARCONI server (CINECA, Bologna, Italy).

## Notes

### Competing Interest Statement

The authors have declared no competing interest.

