## Supplementary Material 1 for "Diel transcriptional pattern contributes to functional and taxonomic diversity in supraglacial microbial communities": coxA.krona.taxa.4_30.html

Javascript must be enabled to view this page.

members
magnitude
magnitudeUnassigned
count
unassigned
taxon
rank
score

coxA.gene2kegg2abundance2tax.norm.forni.trans


1

A09100 Metabolism
71
0
898.965410968644

2759
190
1
7.80544407253
superkingdom

1
190
4751
kingdom
7.80544407253

451864
190
1
subkingdom
7.80544407253

7.80544407253
phylum
1
190
5204

subphylum
7.80544407253
1
190
452284

class
7.80544407253
1
190
1538075

order
7.80544407253
162474
190
1

742845
190
1
7.80544407253
family

genus
7.80544407253
55193
190
1


A09100 Metabolism
species
7.80544407253
1
190
76777

0

A09100 MetabolismA09100 MetabolismA09100 Metabolism
891.159966896114
superkingdom
69
3
190
2

12.0166667729
phylum
190
1297
2

2
188787
190
class
12.0166667729

68933
190
2
12.0166667729
order

188786
190
2
family
12.0166667729

270
190
2
1
12.0166667729
genus

A09100 Metabolism
12.0166667729

0
species

A09100 Metabolism
88190
0
1

31
190
1224
256.214818266649
phylum

2
68525
0
0
subphylum

2
0
28221
class
0

1797802
0
1
species
0

A09100 Metabolism

order
0
0
29
1

1
0
80811
0
suborder

1
0
1524215
family
0

1
0
161492

A09100 Metabolism
genus
0

20.760287774
class
190
1236
4

order
0
0

A09100 Metabolism
0
72274
1
2

family
0
0
135621
1

286
0
1
0
genus

A09100 Metabolism

135614
0
1
0
order

0
32033
1
family
0

0
83614
1
0
genus

1
0
83615

A09100 Metabolism
0
species

order
20.760287774
190
135613
1

1
190
72276

A09100 Metabolism
20.760287774
family

10
1
28216
190
0

A09100 Metabolism
81.768153982089
class

1
206389
190
0.497261240005
order

75787
190
1
family
0.497261240005

1
190
1914449
0.497261240005
genus

1
190
1565605

A09100 Metabolism
species
0.497261240005

19.704525165384

A09100 MetabolismA09100 Metabolism
81.270892742084
order
8
2
80840
190

0
316612
1
0
genus

1
1500261
0

A09100 Metabolism
0
species

genus
0
1
0
212743

0
species

A09100 Metabolism
1736528
0
1

family
61.5663675767
4
190
80864


A09100 MetabolismA09100 Metabolism
genus
33.8632263556
2
80865
190

1
190
34072

A09100 Metabolism
27.7031412211
genus

0
genus
1
0
335058

1
1736430
0

A09100 Metabolism
species
0

190
28211
15
class
153.68637651056

7
2
356
190

A09100 MetabolismA09100 Metabolism
0.615075267505
1.42948425796
order

family
0
0
45401
2

1
2
81
0

A09100 Metabolism
0
genus
0

species
0

A09100 Metabolism
121290
0
1

family
0.814408990455
335928
190
3

genus
0
1
0
152053

1
0
921

A09100 Metabolism
0
species


A09100 Metabolism
genus
0.608764274515
1
279
190


A09100 Metabolism
0.20564471594
genus
1
190
99

4
204455
190
order
152.2568922526

31989
190
4
family
152.2568922526

152.2568922526
genus
121.665156259

A09100 MetabolismA09100 MetabolismA09100 Metabolism
265
190
4
3

1
190
147645

A09100 Metabolism
species
30.5917359936

0
204457
3
order
0

3
1
41297
0
0

A09100 Metabolism
0
family

1
13687
0
genus
0


A09100 Metabolism
species
0
1
0
1380389

0
genus

A09100 Metabolism
362865
0
1

204458
0
1
0
order

0
family
0
76892
1

genus
0
1
75
0

species
0

A09100 Metabolism
88688
0
1

2
0
74201
phylum
0

class
0
1
203494
0

48461
0
1
order
0


A09100 Metabolism
family
0
1
134627
0

0
class
1
414999
0

1
415000
0
order
0

0
family
1
0
134623

0
genus
0
178440
1

0
species

A09100 Metabolism
0
107709
1


A09100 MetabolismA09100 MetabolismA09100 Metabolism
7.69070777103
phylum
19.91871172923
3
6
1117
190

subclass
12.2280039582
2
190
1301283

190
1150
2
12.2280039582
order

A09100 MetabolismA09100 Metabolism

0
order
0
1890424
1

1
0
1890438
0
family

0
genus
1
0
47251

1752064
0
1
species
0

A09100 Metabolism


A09100 Metabolism
0
0
phylum
2
1
0
976

0
class
1853228
0
1

0
order
0
1853229
1

0
family

A09100 Metabolism
563835
0
1

23
201174
190
phylum
603.009770127335

0

A09100 MetabolismA09100 MetabolismA09100 Metabolism
603.009770127335
class
23
3
1760
190

0
order
1
85011
0

2062
0
1
family
0


A09100 Metabolism
0
genus
1
1883
0

0
order
1
0
85008

1
0
28056

A09100 Metabolism
family
0

190
85009
4
562.58809234515
order

3
4
190
31957
260.39941400415

A09100 MetabolismA09100 MetabolismA09100 Metabolism
family
562.58809234515

1
190
1912215
genus
302.188678341

302.188678341
species

A09100 Metabolism
190
1751
1

order
0
0

A09100 Metabolism
0
85006
1
7

0
family
85021
0
2

genus
0
53357
0
2

0
1386088
2
0
species

A09100 MetabolismA09100 Metabolism

0
85023
4
2
0
family

A09100 MetabolismA09100 Metabolism
0

0

A09100 Metabolism
0
genus
2
1
447237
0


A09100 Metabolism
species
0
1
1736329
0


A09100 Metabolism
0
order
40.4216777821851
1
7
85007
190

1
85026
190
17.3852799182
family

genus
17.3852799182
2053
190
1

1
2054
190

A09100 Metabolism
17.3852799182
species

190
1653
1
5
family
23.0363978639851

A09100 Metabolism
0.0979107374951

190
1716
4
22.93848712649
genus

A09100 MetabolismA09100 MetabolismA09100 MetabolismA09100 Metabolism
