## Supplementary Material 1 for "Diel transcriptional pattern contributes to functional and taxonomic diversity in supraglacial microbial communities": coxA.krona.taxa.7_30.html

Javascript must be enabled to view this page.

members
magnitude
magnitudeUnassigned
count
unassigned
taxon
rank
score

coxA.gene2kegg2abundance2tax.norm.forni.trans


A09100 Metabolism
71
283.408552540681
1
0

superkingdom

A09100 MetabolismA09100 MetabolismA09100 Metabolism
190
10.38789623237
2
3
69
283.408552540681

phylum
190
1297
11.02408995848
2

188787
11.02408995848
2
class
190

68933
2
11.02408995848
order
190

11.02408995848
2
188786
190
family

1.51096596619
270
1
2
11.02408995848
genus

A09100 Metabolism
190

1
9.51312399229
88190
190

A09100 Metabolism
species

190
phylum
31
54.773672923791
1224

190
class
0.556838601235
4
1236

190
order
1
0.556838601235
135613

family
190

A09100 Metabolism
72276
1
0.556838601235

0

A09100 Metabolism
order
1
2
0
0
72274

family
0
135621
0
1


A09100 Metabolism
0
genus
1
0
286

order
0
135614
1
0

0
1
32033
0
family

0
1
83614
0
genus

83615
1
0
species
0

A09100 Metabolism

190
subphylum
2
9.86542753868
68525

9.86542753868
2
28221
190
class

species
190

A09100 Metabolism
1797802
6.07145032025
1

190
order
1
3.79397721843
29

suborder
190
80811
1
3.79397721843

family
190
1524215
1
3.79397721843


A09100 Metabolism
190
genus
3.79397721843
1
161492

7.49653845107
28216
10
21.693477548366
1
class

A09100 Metabolism
190

190
order
1
1.71294679686
206389

family
190
75787
1
1.71294679686

genus
190
1914449
1
1.71294679686

1565605
1
1.71294679686
species

A09100 Metabolism
190

0
80840
2
12.483992300436
8
order
190

A09100 MetabolismA09100 Metabolism

0
1
316612
0
genus

species
0

A09100 Metabolism
1500261
0
1

4
12.206235804827
80864
190
family

genus

A09100 Metabolism
190
34072
2.12140364245
1


A09100 MetabolismA09100 Metabolism
190
genus
9.65616993296
2
80865

190
genus
0.428662229417
1
335058

1736430
0.428662229417
1
species

A09100 Metabolism
190

0.277756495609
1
212743
190
genus

1
0.277756495609
1736528

A09100 Metabolism
190
species

190
class
22.65792923551
15
28211

2
7
8.70990444707
3.23696026997
356

A09100 MetabolismA09100 Metabolism
190
order

2
1.33474640019
45401
190
family

genus
190

A09100 Metabolism
1.33474640019
81
2
1.33474640019
1

121290
0
1
species

A09100 Metabolism
0

335928
4.13819777691
3
family
190

279
1
0
genus
0

A09100 Metabolism

genus
190

A09100 Metabolism
99
1
2.23293844588

1
1.90525933103
152053
190
genus

1
1.90525933103
921

A09100 Metabolism
190
species

3
13.94802478844
204457
190
order

41297
2.89988402984
1
3
13.94802478844
family
190

A09100 Metabolism

1
0
13687
0
genus

species

A09100 Metabolism
0
1380389
1
0

genus

A09100 Metabolism
190
362865
11.0481407586
1

0
order
0
1
204458

family
0
76892
0
1

0
1
75
0
genus

0
1
88688
0

A09100 Metabolism
species

order
0
204455
4
0

0
family
4
0
31989


A09100 MetabolismA09100 MetabolismA09100 Metabolism
0
genus
0
4
3
0
265


A09100 Metabolism
0
species
0
1
147645

976
0
1
0
2
phylum

A09100 Metabolism
0

0
class
1
0
1853228

1
0
1853229
0
order

1
0
563835
0

A09100 Metabolism
family

23
128.41748474012
201174
190
phylum

10.61170113039
1760
3
23
128.41748474012
class
190

A09100 MetabolismA09100 MetabolismA09100 Metabolism

order
190

A09100 Metabolism
85007
6.79577990351
18.41469433453
7
1

0
family
0
1
85026

2053
1
0
genus
0

2054
0
1
species

A09100 Metabolism
0

1
5
11.61891443102
1653
1.24250647112
190

A09100 Metabolism
family

1716
10.3764079599
4
genus

A09100 MetabolismA09100 MetabolismA09100 MetabolismA09100 Metabolism
190

56.9732062059
4
85009
190
order

31957
24.660445163
3
4
56.9732062059
family

A09100 MetabolismA09100 MetabolismA09100 Metabolism
190

genus
190
1912215
1
32.3127610429


A09100 Metabolism
190
species
32.3127610429
1
1751

order

A09100 Metabolism
190
85006
0
1
7
35.19973315135

25.527014474
2
85021
190
family

genus
190
53357
2
25.527014474

2
25.527014474
1386088

A09100 MetabolismA09100 Metabolism
190
species

family

A09100 MetabolismA09100 Metabolism
190
85023
3.75540832904
2
9.67271867735
4

447237
5.91731034831
5.91731034831
2
1
genus
190

A09100 Metabolism

1736329
1
0
species
0

A09100 Metabolism

order
190
85008
7.21814991795
1

190

A09100 Metabolism
family
1
7.21814991795
28056

0
order
0
1
85011

2062
1
0
family
0

1
0
1883
0

A09100 Metabolism
genus

26.3284296877
2
74201
190
phylum

414999
12.4367459324
1
class
190

order
190
415000
12.4367459324
1

190
family
1
12.4367459324
134623

178440
1
12.4367459324
genus
190

190

A09100 Metabolism
species
1
12.4367459324
107709

class
190
203494
1
13.8916837553

190
order
1
13.8916837553
48461

1
13.8916837553
134627
190

A09100 Metabolism
family

phylum
190

A09100 MetabolismA09100 MetabolismA09100 Metabolism
1117
28.36542607716
6
52.47697899822
3

order
190
1890424
15.5688776715
1

family
190
1890438
15.5688776715
1

genus
190
47251
1
15.5688776715

species
190

A09100 Metabolism
1752064
1
15.5688776715

190
subclass
8.54267524956
2
1301283

order
190

A09100 MetabolismA09100 Metabolism
1150
8.54267524956
2

2759
0
1
superkingdom
0

4751
0
1
kingdom
0

subkingdom
0
451864
0
1

phylum
0
5204
1
0

subphylum
0
452284
1
0

1
0
1538075
0
class

0
order
1
0
162474

0
family
0
1
742845

genus
0
55193
1
0

76777
1
0
species

A09100 Metabolism
0
