## Supplementary Material 1 for "Diel transcriptional pattern contributes to functional and taxonomic diversity in supraglacial microbial communities": coxA.krona.taxa.13_30.html

Javascript must be enabled to view this page.

members
magnitude
magnitudeUnassigned
count
unassigned
taxon
rank
score

coxA.gene2kegg2abundance2tax.norm.forni.trans


A09100 Metabolism
71
677.635627296126
1
0

3
0
644.472779014926

A09100 MetabolismA09100 MetabolismA09100 Metabolism
190
2
superkingdom
69

1117
phylum
6
0
3
0

A09100 MetabolismA09100 MetabolismA09100 Metabolism
0

0
0
2
1301283
subclass

order
1150
2
0
0

A09100 MetabolismA09100 Metabolism

0
0
1
1890424
order

1
1890438
family
0
0

1
47251
genus
0
0

0

A09100 Metabolism
0
1
species
1752064

31
phylum
1224
190
163.37850482762

10
class
28216

A09100 Metabolism
190
88.90009656445
0
1

87.6388910265
26.6533887328
2

A09100 MetabolismA09100 Metabolism
190
order
80840
8

316612
genus
1
0
0

species
1500261
1
0
0

A09100 Metabolism

family
80864
4
60.9855022937
190

0
0

A09100 Metabolism
genus
34072
1

335058
genus
1
0
0


A09100 Metabolism
0
0
1
1736430
species

60.9855022937

A09100 MetabolismA09100 Metabolism
190
80865
genus
2

0
0
1
genus
212743

0
0

A09100 Metabolism
1736528
species
1

order
206389
1
1.26120553795
190

family
75787
1
1.26120553795
190

1.26120553795
190
1914449
genus
1

1
species
1565605
190

A09100 Metabolism
1.26120553795

class
1236
4
68.0432056623
190

2
order
72274
190

A09100 Metabolism
35.7333170744
35.7333170744
1

family
135621
1
0
0

0

A09100 Metabolism
0
1
286
genus

order
135614
1
32.3098885879
190

32.3098885879
190
family
32033
1

32.3098885879
190
genus
83614
1

32.3098885879

A09100 Metabolism
190
species
83615
1

135613
order
1
0
0

72276
family
1
0
0

A09100 Metabolism

0
0
68525
subphylum
2

2
class
28221
0
0

species
1797802
1
0
0

A09100 Metabolism

0
0
1
29
order

1
80811
suborder
0
0

family
1524215
1
0
0

0

A09100 Metabolism
0
1
genus
161492

6.43520260087
190
class
28211
15

0.35766574512
190
order
204455
4

4
31989
family
190
0.35766574512

0.35766574512
0.35766574512
3
190

A09100 MetabolismA09100 MetabolismA09100 Metabolism
genus
265
4


A09100 Metabolism
0
0
1
147645
species

1
204458
order
0
0

76892
family
1
0
0

genus
75
1
0
0

1
88688
species
0

A09100 Metabolism
0

7
356
order
190

A09100 MetabolismA09100 Metabolism
2
1.536113532895
6.07753685575

190
2.199027126278
3
family
335928

genus
99
1
1.69078077384

A09100 Metabolism
190

190
0.508246352438
1
genus
152053

species
921
1
0.508246352438

A09100 Metabolism
190

279
genus
1
0
0

A09100 Metabolism

family
45401
2
2.342396196577
190


A09100 Metabolism
190
2.342396196577
1.58754594421
1
2
genus
81

0.754850252367
190

A09100 Metabolism
121290
species
1

204457
order
3
0
0

3
family
41297

A09100 Metabolism
0
0
0
1

genus
362865
1
0
0

A09100 Metabolism

genus
13687
1
0
0

1380389
species
1
0

A09100 Metabolism
0

201174
phylum
23
481.094274187306
190

481.094274187306
3
0.276586703699
190

A09100 MetabolismA09100 MetabolismA09100 Metabolism
class
1760
23

7
order
85006
190

A09100 Metabolism
0.211158094122
0
1

0
2
0.211158094122
190

A09100 MetabolismA09100 Metabolism
85023
family
4

genus
447237
2
0.211158094122
0
1
190

A09100 Metabolism

0.211158094122
190

A09100 Metabolism
1736329
species
1

0
0
family
85021
2

2
genus
53357
0
0

0

A09100 MetabolismA09100 Metabolism
0
1386088
species
2

28.260079299416
0
1
190

A09100 Metabolism
order
85007
7

1
0
28.260079299416

A09100 Metabolism
190
1653
family
5

4
genus
1716
190

A09100 MetabolismA09100 MetabolismA09100 MetabolismA09100 Metabolism
28.260079299416

1
85026
family
0
0

0
0
1
genus
2053

2054
species
1
0

A09100 Metabolism
0

452.346450090069
190
85009
order
4

4
31957
family

A09100 MetabolismA09100 MetabolismA09100 Metabolism
190
172.010006293069
3
452.346450090069

280.336443797
190
genus
1912215
1

1
1751
species
190

A09100 Metabolism
280.336443797

order
85008
1
0
0

0

A09100 Metabolism
0
family
28056
1

85011
order
1
0
0

2062
family
1
0
0

0

A09100 Metabolism
0
1
1883
genus

0
0
74201
phylum
2

1
class
414999
0
0

1
order
415000
0
0

1
family
134623
0
0

178440
genus
1
0
0

0

A09100 Metabolism
0
species
107709
1

1
class
203494
0
0

1
order
48461
0
0

1
family
134627
0

A09100 Metabolism
0

0
0
2
1297
phylum

2
class
188787
0
0

0
0
2
68933
order

2
188786
family
0
0

1
0
0

A09100 Metabolism
0
270
genus
2


A09100 Metabolism
0
0
1
species
88190

0
1
0

A09100 Metabolism
0
phylum
976
2

0
0
class
1853228
1

0
0
1
order
1853229

1
family
563835
0

A09100 Metabolism
0

33.1628482812
190
2759
superkingdom
1

kingdom
4751
1
33.1628482812
190

33.1628482812
190
451864
subkingdom
1

1
phylum
5204
190
33.1628482812

1
452284
subphylum
190
33.1628482812

1
class
1538075
190
33.1628482812

190
33.1628482812
1
order
162474

1
742845
family
190
33.1628482812

33.1628482812
190
genus
55193
1

76777
species
1
33.1628482812

A09100 Metabolism
190
