## Supplementary Material 1 for "Diel transcriptional pattern contributes to functional and taxonomic diversity in supraglacial microbial communities": coxA.krona.taxa.19_30.html

Javascript must be enabled to view this page.

members
magnitude
magnitudeUnassigned
count
unassigned
taxon
rank
score

coxA.gene2kegg2abundance2tax.norm.forni.trans


1
1221.865175
71
6.35973

A09100 Metabolism


A09100 MetabolismA09100 MetabolismA09100 Metabolism
2
superkingdom
69
190
82.5725
3
1188.302045

19.2932
190
29.6885
1
2

A09100 Metabolism
976
phylum

class
1853228
1
190
10.3953

1
order
1853229
10.3953
190

10.3953
190
1
family
563835

A09100 Metabolism

190
15.471874
phylum
1297
2

15.471874
190
2
class
188787

2
order
68933
190
15.471874

family
188786
2
15.471874
190

2
genus
270

A09100 Metabolism
190
0.483974
1
15.471874

14.9879
190
1

A09100 Metabolism
species
88190

688.44958
190
23
201174
phylum

60.5687
190
688.44958
3

A09100 MetabolismA09100 MetabolismA09100 Metabolism
1760
class
23

190
410.3072
4
85009
order

4

A09100 MetabolismA09100 MetabolismA09100 Metabolism
family
31957
410.3072
3
211.5702
190

190
198.737
genus
1912215
1

190
198.737
1
1751

A09100 Metabolism
species

0
0
1
order
85008

0
0
1
family
28056

A09100 Metabolism

7

A09100 Metabolism
order
85006
15.9397
190
125.22685
1

2
85021
family
13.90285
190

53357
genus
2
190
13.90285


A09100 MetabolismA09100 Metabolism
species
1386088
2
13.90285
190

95.3843
2
36.4134
190
4

A09100 MetabolismA09100 Metabolism
85023
family

190
14.2543
1
58.9709
447237

A09100 Metabolism
genus
2

190
44.7166
1

A09100 Metabolism
species
1736329

190
7.70316
1
62.46803
7
85007

A09100 Metabolism
order

85026
family
1
6.01866
190

190
6.01866
1
genus
2053

2054
species

A09100 Metabolism
1
6.01866
190

family
1653

A09100 Metabolism
5
1
48.74621
190
23.7929

190
24.95331
4

A09100 MetabolismA09100 MetabolismA09100 MetabolismA09100 Metabolism
genus
1716

order
85011
1
190
29.8788

1
family
2062
29.8788
190

190
29.8788
1
1883
genus

A09100 Metabolism

190
43.64124
3
67.613384
6
phylum
1117

A09100 MetabolismA09100 MetabolismA09100 Metabolism

2
subclass
1301283
190
23.60741

23.60741
190
order

A09100 MetabolismA09100 Metabolism
1150
2

1
1890424
order
190
0.364734

1
family
1890438
0.364734
190

0.364734
190
1
genus
47251

190
0.364734

A09100 Metabolism
species
1752064
1

190
285.720897
31
phylum
1224

5.7411
190
157.30477
1
10

A09100 Metabolism
class
28216

206389
order
1
190
1.78072

190
1.78072
75787
family
1

1
1914449
genus
1.78072
190

1.78072
190
1
1565605
species

A09100 Metabolism

2
149.78295
190
14.00492
8
80840

A09100 MetabolismA09100 Metabolism
order

190
44.92063
80864
family
4

190
25.16983
2
genus
80865

A09100 MetabolismA09100 Metabolism

0
0
1
34072

A09100 Metabolism
genus

1
335058
genus
190
19.7508


A09100 Metabolism
1736430
species
1
190
19.7508

1
212743
genus
46.2875
190

190
46.2875
1736528
species

A09100 Metabolism
1

1
316612
genus
44.5699
190

species
1500261

A09100 Metabolism
1
44.5699
190

190
6.2005
68525
subphylum
2

2
28221
class
6.2005
190

29
order
1
190
6.2005

6.2005
190
1
suborder
80811

1
family
1524215
190
6.2005

190
6.2005

A09100 Metabolism
genus
161492
1


A09100 Metabolism
species
1797802
1
0
0

79.008557
190
class
28211
15

1
order
204458
34.108
190

1
76892
family
34.108
190

34.108
190
75
genus
1

species
88688

A09100 Metabolism
1
190
34.108

204455
order
4
190
8.487058

8.487058
190
4
31989
family

265
genus

A09100 MetabolismA09100 MetabolismA09100 Metabolism
4
3
8.487058
190
7.571906

1

A09100 Metabolism
147645
species
190
0.915152

356
order

A09100 MetabolismA09100 Metabolism
7
3.119109
2
0.63189
190

3
335928
family
190
1.449029

279
genus

A09100 Metabolism
1
0
0

190
0.792651
1
152053
genus

0.792651
190

A09100 Metabolism
921
species
1

190
0.656378
1
genus
99

A09100 Metabolism

1.03819
190
2
45401
family

2

A09100 Metabolism
genus
81
0
190
1.03819
1

species

A09100 Metabolism
121290
1
1.03819
190

order
204457
3
33.29439
190

family

A09100 Metabolism
41297
3
33.29439
1
11.1085
190

1
genus
13687
190
13.6701

13.6701
190
1380389
species

A09100 Metabolism
1

1
362865

A09100 Metabolism
genus
190
8.51579

43.20707
190
1236
class
4


A09100 Metabolism
order
72274
2
1
25.5649
190
12.4515

135621
family
1
13.1134
190

13.1134
190
1

A09100 Metabolism
286
genus

order
135613
1
190
13.1554

1
family
72276

A09100 Metabolism
190
13.1554

4.48677
190
135614
order
1

4.48677
190
1
family
32033

4.48677
190
1
genus
83614


A09100 Metabolism
83615
species
1
4.48677
190

18.78531
190
phylum
74201
2

15.4742
190
414999
class
1

order
415000
1
15.4742
190

1
family
134623
15.4742
190

genus
178440
1
190
15.4742


A09100 Metabolism
107709
species
1
15.4742
190

190
3.31111
1
class
203494

3.31111
190
48461
order
1

3.31111
190

A09100 Metabolism
family
134627
1

27.2034
190
1
2759
superkingdom

1
4751
kingdom
27.2034
190

190
27.2034
1
451864
subkingdom

190
27.2034
5204
phylum
1

190
27.2034
452284
subphylum
1

1
class
1538075
27.2034
190

190
27.2034
1
order
162474

742845
family
1
190
27.2034

27.2034
190
1
55193
genus

1
species

A09100 Metabolism
76777
27.2034
190
