## Supplementary Material 1 for "Diel transcriptional pattern contributes to functional and taxonomic diversity in supraglacial microbial communities": narG.krona.taxa.4_30.html

Javascript must be enabled to view this page.

members
magnitude
magnitudeUnassigned
count
unassigned
taxon
rank
score

narG.gene2kegg2abundance2tax.norm.forni.trans


1495.92193044936
92

10239
0
2
superkingdom
0

1608440
0

A09100 MetabolismA09130 Environmental Information Processing
species
2
0

2

A09100 MetabolismA09130 Environmental Information ProcessingA09100 MetabolismA09130 Environmental Information ProcessingA09100 MetabolismA09130 Environmental Information ProcessingA09100 MetabolismA09130 Environmental Information Processing
1495.92193044936
90
8
superkingdom
181.191272795648
1465


A09100 MetabolismA09130 Environmental Information Processing
0
77133
0
2
species

2
phylum
0
1297
0

2
class
0
188787
0

68933
0
2
order
0

2
family
0
188786
0

0

A09100 MetabolismA09130 Environmental Information Processing
270
0
genus
2

961.783950705736
201174
1465
24
phylum

1465
class
10
24
0

A09100 MetabolismA09130 Environmental Information ProcessingA09100 MetabolismA09130 Environmental Information ProcessingA09100 MetabolismA09130 Environmental Information ProcessingA09100 MetabolismA09130 Environmental Information ProcessingA09100 MetabolismA09130 Environmental Information Processing
961.783950705736
1760

961.3541909466
85009
1465
order
12


A09100 MetabolismA09130 Environmental Information ProcessingA09100 MetabolismA09130 Environmental Information ProcessingA09100 MetabolismA09130 Environmental Information Processing
961.3541909466
31957
1465
12
family
6
716.4903104414

1912216
21.1605245292
genus
4
1465

4
species
1465
1050843
21.1605245292

A09100 MetabolismA09130 Environmental Information ProcessingA09100 MetabolismA09130 Environmental Information Processing

1465
2
genus
223.703355976
1912215

species
2
1465
1748
223.703355976

A09100 MetabolismA09130 Environmental Information Processing

1465
2
order
0.429759759136
2037

0.429759759136
2049
1465
family
2


A09100 MetabolismA09130 Environmental Information Processing
0.429759759136
1654
1465
2
genus

1465
phylum
14
85.2411625905
1239

1465
class
14
85.2411625905
91061

1465
order
14
85.2411625905
1385

85.2411625905
90964
1465
family
14

1279

A09100 MetabolismA09130 Environmental Information ProcessingA09100 MetabolismA09130 Environmental Information ProcessingA09100 MetabolismA09130 Environmental Information ProcessingA09100 MetabolismA09130 Environmental Information ProcessingA09100 MetabolismA09130 Environmental Information ProcessingA09100 MetabolismA09130 Environmental Information ProcessingA09100 MetabolismA09130 Environmental Information Processing
85.2411625905
14
genus
1465

1465
phylum
40
267.705544357473
1224


A09100 MetabolismA09130 Environmental Information Processing
62.7725773747126
28216
1465
class
2
26
1.470838636306

80840
60.9600777277172

A09100 MetabolismA09130 Environmental Information Processing
0
order
2
20
1465

0
genus
8
0
212743

0

A09100 MetabolismA09130 Environmental Information ProcessingA09100 MetabolismA09130 Environmental Information ProcessingA09100 MetabolismA09130 Environmental Information ProcessingA09100 MetabolismA09130 Environmental Information Processing
1736528
0
8
species

60.9600777277172
80864
1465
family
10

8
genus
1465
80865
60.877701564

A09100 MetabolismA09130 Environmental Information ProcessingA09100 MetabolismA09130 Environmental Information ProcessingA09100 MetabolismA09130 Environmental Information ProcessingA09100 MetabolismA09130 Environmental Information Processing

0.0823761637172
34072
1465
2
genus


A09100 MetabolismA09130 Environmental Information Processing
0.0823761637172
1882774
1465
species
2

1465
4
order
0.3416610106894
32003

1465
4
family
0.3416610106894
2008793

0.3416610106894
378210
1465
4
genus

4
species
1465
1842540

A09100 MetabolismA09130 Environmental Information ProcessingA09100 MetabolismA09130 Environmental Information Processing
0.3416610106894

196.09350625594

A09100 MetabolismA09130 Environmental Information Processing
28211
1465
56.1694923022
2
class
8

0
2
order
0

A09100 MetabolismA09130 Environmental Information Processing
356

204455
139.92401395374
order
4
1465

4
family
1465
31989
139.92401395374

17.47254834254
265
1465
genus
2

147645

A09100 MetabolismA09130 Environmental Information Processing
17.47254834254
species
2
1465

122.4514656112
1400060
1465
2
genus

species
2
1465
1666912
122.4514656112

A09100 MetabolismA09130 Environmental Information Processing

1236

A09100 MetabolismA09130 Environmental Information Processing
8.83946072682
6
class
2
4.93994879186
1465

3.89951193496
order
4
2
1465
91347
3.89951193496

A09100 MetabolismA09130 Environmental Information Processing

0
family
2
0
1903411

0
613
0
2
genus

species
2
0
614

A09100 MetabolismA09130 Environmental Information Processing
0
