## Supplementary Material 1 for "Diel transcriptional pattern contributes to functional and taxonomic diversity in supraglacial microbial communities": narG.krona.taxa.7_30.html

Javascript must be enabled to view this page.

members
magnitude
magnitudeUnassigned
count
unassigned
taxon
rank
score

narG.gene2kegg2abundance2tax.norm.forni.trans


291.57073808293
92


A09100 MetabolismA09130 Environmental Information ProcessingA09100 MetabolismA09130 Environmental Information ProcessingA09100 MetabolismA09130 Environmental Information ProcessingA09100 MetabolismA09130 Environmental Information Processing
1465
8
254.41444209413
superkingdom
2
54.92902377952
90

2
1297
1.083725299676
phylum
1465

class
1.083725299676
1465
2
188787

1.083725299676
order
1465
2
68933

188786
2
family
1.083725299676
1465

genus
1.083725299676

A09100 MetabolismA09130 Environmental Information Processing
1465
2
270

1239
14
1465
phylum
11.1607289579

91061
14
1465
class
11.1607289579

order
11.1607289579
1465
14
1385

1465
family
11.1607289579
90964
14

1279
14

A09100 MetabolismA09130 Environmental Information ProcessingA09100 MetabolismA09130 Environmental Information ProcessingA09100 MetabolismA09130 Environmental Information ProcessingA09100 MetabolismA09130 Environmental Information ProcessingA09100 MetabolismA09130 Environmental Information ProcessingA09100 MetabolismA09130 Environmental Information ProcessingA09100 MetabolismA09130 Environmental Information Processing
1465
11.1607289579
genus

1465
43.248064333308
phylum
40
1224


A09100 MetabolismA09130 Environmental Information Processing
1465
2
class
43.248064333308
5.62457060054
28216
26

1465
order
5.37709558504
32003
4

4
2008793
1465
family
5.37709558504

378210
4
1465
5.37709558504
genus

1842540
4
5.37709558504
species

A09100 MetabolismA09130 Environmental Information ProcessingA09100 MetabolismA09130 Environmental Information Processing
1465


A09100 MetabolismA09130 Environmental Information Processing
1465
2
order
32.246398147728
0
80840
20

212743
8
1465
16.44854663782
genus

1465

A09100 MetabolismA09130 Environmental Information ProcessingA09100 MetabolismA09130 Environmental Information ProcessingA09100 MetabolismA09130 Environmental Information ProcessingA09100 MetabolismA09130 Environmental Information Processing
species
16.44854663782
1736528
8

1465
15.797851509908
family
80864
10

genus
14.85421638474

A09100 MetabolismA09130 Environmental Information ProcessingA09100 MetabolismA09130 Environmental Information ProcessingA09100 MetabolismA09130 Environmental Information ProcessingA09100 MetabolismA09130 Environmental Information Processing
1465
80865
8

34072
2
0.943635125168
genus
1465

1465

A09100 MetabolismA09130 Environmental Information Processing
0.943635125168
species
2
1882774

6
1236
0
class
0
2

A09100 MetabolismA09130 Environmental Information Processing
0

4
0
91347
0

A09100 MetabolismA09130 Environmental Information Processing
order
0
2

1903411
2
0
family
0

0
genus
0
2
613

0

A09100 MetabolismA09130 Environmental Information Processing
species
0
2
614

0
28211
8
2
class
0
0

A09100 MetabolismA09130 Environmental Information Processing

0
0
order
4
204455

31989
4
0
0
family

2
265
0
genus
0

species
0

A09100 MetabolismA09130 Environmental Information Processing
0
147645
2

0
genus
0
2
1400060

0

A09100 MetabolismA09130 Environmental Information Processing
0
species
2
1666912

356
2
0
order

A09100 MetabolismA09130 Environmental Information Processing
0

24
201174
142.96714457034
phylum
1465

142.96714457034
class
10

A09100 MetabolismA09130 Environmental Information ProcessingA09100 MetabolismA09130 Environmental Information ProcessingA09100 MetabolismA09130 Environmental Information ProcessingA09100 MetabolismA09130 Environmental Information ProcessingA09100 MetabolismA09130 Environmental Information Processing
1465
24
1760
9.74189862416

85009
12
133.22524594618
order
1465

6
133.22524594618
family
1465

A09100 MetabolismA09130 Environmental Information ProcessingA09100 MetabolismA09130 Environmental Information ProcessingA09100 MetabolismA09130 Environmental Information Processing
99.880282133
31957
12

1465
genus
12.91346491458
4
1912216

4
1050843

A09100 MetabolismA09130 Environmental Information ProcessingA09100 MetabolismA09130 Environmental Information Processing
1465
species
12.91346491458

20.4314988986
genus
1465
1912215
2

1748
2
1465

A09100 MetabolismA09130 Environmental Information Processing
20.4314988986
species

2037
2
0
order
0

0
family
0
2
2049

0
genus
0

A09100 MetabolismA09130 Environmental Information Processing
1654
2

2
77133

A09100 MetabolismA09130 Environmental Information Processing
1465
1.025755153386
species

10239
2
superkingdom
37.1562959888
1465

1465

A09100 MetabolismA09130 Environmental Information Processing
species
37.1562959888
2
1608440
