## Supplementary Material 1 for "Diel transcriptional pattern contributes to functional and taxonomic diversity in supraglacial microbial communities": narG.krona.taxa.13_30.html

Javascript must be enabled to view this page.

members
magnitude
magnitudeUnassigned
count
unassigned
taxon
rank
score

narG.gene2kegg2abundance2tax.norm.forni.trans


1097.14597327908
92

2

A09100 MetabolismA09130 Environmental Information ProcessingA09100 MetabolismA09130 Environmental Information ProcessingA09100 MetabolismA09130 Environmental Information ProcessingA09100 MetabolismA09130 Environmental Information Processing
8
1086.4750868666
superkingdom
1465
90
120.106738045952


A09100 MetabolismA09130 Environmental Information Processing
77133
0
species
0
2

201174
phylum
1465
808.039523098612
24

24
0.343778245412
1760

A09100 MetabolismA09130 Environmental Information ProcessingA09100 MetabolismA09130 Environmental Information ProcessingA09100 MetabolismA09130 Environmental Information ProcessingA09100 MetabolismA09130 Environmental Information ProcessingA09100 MetabolismA09130 Environmental Information Processing
10
808.039523098612
1465
class

2037
order
0
0
2

2
2049
family
0
0

2
0
0
genus
1654

A09100 MetabolismA09130 Environmental Information Processing

12
1465
order
807.6957448532
85009

598.1810864472
12
family
1465
807.6957448532

A09100 MetabolismA09130 Environmental Information ProcessingA09100 MetabolismA09130 Environmental Information ProcessingA09100 MetabolismA09130 Environmental Information Processing
6
31957

0
genus
0
1912216
4

0
species
0
1050843

A09100 MetabolismA09130 Environmental Information ProcessingA09100 MetabolismA09130 Environmental Information Processing
4

1912215
209.514658406
genus
1465
2

2

A09100 MetabolismA09130 Environmental Information Processing
1748
1465
species
209.514658406

phylum
1465
73.7370934279744
1224
40

21.9219063526
class
1465
1236
2

A09100 MetabolismA09130 Environmental Information Processing
6
21.9219063526

order
0
0
2

A09100 MetabolismA09130 Environmental Information Processing
91347
0
4

1903411
0
family
0
2

613
0
0
genus
2

2

A09100 MetabolismA09130 Environmental Information Processing
614
species
0
0

8
0.251601153866
28211

A09100 MetabolismA09130 Environmental Information Processing
2
14.3043074710324
1465
class

2
1465
order
14.02369683334

A09100 MetabolismA09130 Environmental Information Processing
356

order
1465
0.0290094838264
204455
4

31989
1465
family
0.0290094838264
4

1400060
0.0290094838264
1465
genus
2

2
1666912

A09100 MetabolismA09130 Environmental Information Processing
0.0290094838264
species
1465

genus
0
0
265
2

0
species
0

A09100 MetabolismA09130 Environmental Information Processing
147645
2

37.510879604342
1465
class
28216
2

A09100 MetabolismA09130 Environmental Information Processing
0.402777803574
26

0
20
80840
2

A09100 MetabolismA09130 Environmental Information Processing
35.5312169529
order
1465

family
1465
35.5312169529
80864
10

2
34072
1.40076440009
genus
1465

1882774

A09100 MetabolismA09130 Environmental Information Processing
1.40076440009
species
1465
2

1465
genus
34.13045255281

A09100 MetabolismA09130 Environmental Information ProcessingA09100 MetabolismA09130 Environmental Information ProcessingA09100 MetabolismA09130 Environmental Information ProcessingA09100 MetabolismA09130 Environmental Information Processing
80865
8

genus
0
0
212743
8

8
0
species
0
1736528

A09100 MetabolismA09130 Environmental Information ProcessingA09100 MetabolismA09130 Environmental Information ProcessingA09100 MetabolismA09130 Environmental Information ProcessingA09100 MetabolismA09130 Environmental Information Processing

4
1.576884847868
order
1465
32003

4
2008793
1.576884847868
1465
family

4
1.576884847868
1465
genus
378210

1842540

A09100 MetabolismA09130 Environmental Information ProcessingA09100 MetabolismA09130 Environmental Information Processing
1.576884847868
1465
species
4

2
1297
1465
phylum
33.4974382596

2
33.4974382596
1465
class
188787

2
68933
33.4974382596
1465
order

2
1465
family
33.4974382596
188786

2
genus
1465
33.4974382596

A09100 MetabolismA09130 Environmental Information Processing
270

1239
51.09429403446
phylum
1465
14

class
1465
51.09429403446
91061
14

1385
51.09429403446
1465
order
14

14
51.09429403446
1465
family
90964

14
1465
genus
51.09429403446

A09100 MetabolismA09130 Environmental Information ProcessingA09100 MetabolismA09130 Environmental Information ProcessingA09100 MetabolismA09130 Environmental Information ProcessingA09100 MetabolismA09130 Environmental Information ProcessingA09100 MetabolismA09130 Environmental Information ProcessingA09100 MetabolismA09130 Environmental Information ProcessingA09100 MetabolismA09130 Environmental Information Processing
1279

2
10239
10.67088641248
1465
superkingdom

2
1608440

A09100 MetabolismA09130 Environmental Information Processing
10.67088641248
species
1465
