## Supplementary Material 1 for "Diel transcriptional pattern contributes to functional and taxonomic diversity in supraglacial microbial communities": narG.krona.taxa.19_30.html

Javascript must be enabled to view this page.

members
magnitude
magnitudeUnassigned
count
unassigned
taxon
rank
score

narG.gene2kegg2abundance2tax.norm.forni.trans


1508.029786
92

1465
30.8476
superkingdom
2
10239


A09100 MetabolismA09130 Environmental Information Processing
30.8476
1465
species
1608440
2

2
90
8
superkingdom
1477.182186
1465
163.3933

A09100 MetabolismA09130 Environmental Information ProcessingA09100 MetabolismA09130 Environmental Information ProcessingA09100 MetabolismA09130 Environmental Information ProcessingA09100 MetabolismA09130 Environmental Information Processing

phylum
201174
24
887.888609
1465

class
10
24
1760

A09100 MetabolismA09130 Environmental Information ProcessingA09100 MetabolismA09130 Environmental Information ProcessingA09100 MetabolismA09130 Environmental Information ProcessingA09100 MetabolismA09130 Environmental Information ProcessingA09100 MetabolismA09130 Environmental Information Processing
47.730609
1465
887.888609

order
2037
2
0
0

0
0
family
2
2049

0
0

A09100 MetabolismA09130 Environmental Information Processing
2
1654
genus

840.158
1465
85009
12
order

1465
840.158
641.493

A09100 MetabolismA09130 Environmental Information ProcessingA09100 MetabolismA09130 Environmental Information ProcessingA09100 MetabolismA09130 Environmental Information Processing
12
31957
family
6

genus
1912216
4
0
0

species
1050843
4

A09100 MetabolismA09130 Environmental Information ProcessingA09100 MetabolismA09130 Environmental Information Processing
0
0

genus
1912215
2
198.665
1465


A09100 MetabolismA09130 Environmental Information Processing
198.665
1465
species
1748
2

phylum
14
1239
1465
65.60703

1465
65.60703
class
14
91061

1385
14
order
65.60703
1465

65.60703
1465
family
90964
14

65.60703
1465

A09100 MetabolismA09130 Environmental Information ProcessingA09100 MetabolismA09130 Environmental Information ProcessingA09100 MetabolismA09130 Environmental Information ProcessingA09100 MetabolismA09130 Environmental Information ProcessingA09100 MetabolismA09130 Environmental Information ProcessingA09100 MetabolismA09130 Environmental Information ProcessingA09100 MetabolismA09130 Environmental Information Processing
1279
14
genus

1297
2
phylum
16.8943
1465

2
188787
class
1465
16.8943

order
68933
2
16.8943
1465

16.8943
1465
188786
2
family


A09100 MetabolismA09130 Environmental Information Processing
1465
16.8943
genus
2
270

species
77133
2

A09100 MetabolismA09130 Environmental Information Processing
13.3366
1465

phylum
40
1224
1465
330.062347

8
28211
class
2
1465
9.22707
0

A09100 MetabolismA09130 Environmental Information Processing

7.52464
1465

A09100 MetabolismA09130 Environmental Information Processing
356
2
order

4
204455
order
1465
1.70243

4
31989
family
1465
1.70243

genus
1400060
2
1.70243
1465

1.70243
1465

A09100 MetabolismA09130 Environmental Information Processing
1666912
2
species

0
0
2
265
genus

2
147645
species
0
0

A09100 MetabolismA09130 Environmental Information Processing

6
1236
class
2
1465
39.0107

A09100 MetabolismA09130 Environmental Information Processing
0

39.0107
1465

A09100 MetabolismA09130 Environmental Information Processing
14.9973
91347
4
2
order

24.0134
1465
family
1903411
2

genus
613
2
24.0134
1465


A09100 MetabolismA09130 Environmental Information Processing
24.0134
1465
species
614
2

2.64682

A09100 MetabolismA09130 Environmental Information Processing
281.824577
1465
2
class
28216
26

80840
20
2
order
213.646767
1465

A09100 MetabolismA09130 Environmental Information Processing
11.8424

family
80864
10
26.029167
1465

1465
1.44481
2
34072
genus

1465
1.44481

A09100 MetabolismA09130 Environmental Information Processing
2
1882774
species


A09100 MetabolismA09130 Environmental Information ProcessingA09100 MetabolismA09130 Environmental Information ProcessingA09100 MetabolismA09130 Environmental Information ProcessingA09100 MetabolismA09130 Environmental Information Processing
24.584357
1465
genus
80865
8

genus
212743
8
175.7752
1465

1736528
8
species
175.7752
1465

A09100 MetabolismA09130 Environmental Information ProcessingA09100 MetabolismA09130 Environmental Information ProcessingA09100 MetabolismA09130 Environmental Information ProcessingA09100 MetabolismA09130 Environmental Information Processing

1465
65.53099
4
32003
order

65.53099
1465
family
2008793
4

4
378210
genus
1465
65.53099


A09100 MetabolismA09130 Environmental Information ProcessingA09100 MetabolismA09130 Environmental Information Processing
65.53099
1465
species
1842540
4
