## Supplementary Material 1 for "Diel transcriptional pattern contributes to functional and taxonomic diversity in supraglacial microbial communities": psbd.krona.taxa.4_30.html

Javascript must be enabled to view this page.

members
magnitude
magnitudeUnassigned
count
unassigned
taxon
rank
score

psbD.gene2kegg2abundance2tax.norm.forni.trans


2.11094276589
7

0
3
0
2759
superkingdom


A09100 MetabolismA09100 Metabolism
0
0
3
2
0
33090
kingdom

35493
phylum
0
0
1

subphylum
131221
1
0

A09100 Metabolism
0

2.11094276589
4
195
superkingdom
2

phylum
1117
2.11094276589
4
195

1
2.11094276589
195
subclass
1301283

195

A09100 Metabolism
1
2.11094276589
1150
order

order
1890424
0
1
3

A09100 Metabolism
0
0

family
1890438
2
0
0

2
0

A09100 MetabolismA09100 Metabolism
0
genus
47251
