## Supplementary Material 1 for "Diel transcriptional pattern contributes to functional and taxonomic diversity in supraglacial microbial communities": psbd.krona.taxa.7_30.html

Javascript must be enabled to view this page.

members
magnitude
magnitudeUnassigned
count
unassigned
taxon
rank
score

psbD.gene2kegg2abundance2tax.norm.forni.trans


317.43687405351
7

195
310.73600407251
2
superkingdom
4

4
phylum
1117
310.73600407251
195

1301283
195
164.72503571
1
subclass

195
164.72503571
1150
order
1

A09100 Metabolism

195
146.01096836251
4.30294471491
1
1890424
order
3

A09100 Metabolism

1890438
141.7080236476
195
2
family

141.7080236476
195
47251
genus

A09100 MetabolismA09100 Metabolism
2

3
superkingdom
2759
195
6.700869981

33090
195
6.700869981
6.700869981
2
3

A09100 MetabolismA09100 Metabolism
kingdom

1
phylum
35493
0
0

131221
0
0

A09100 Metabolism
1
subphylum
