## Supplementary Material 1 for "Diel transcriptional pattern contributes to functional and taxonomic diversity in supraglacial microbial communities": psbd.krona.taxa.13_30.html

Javascript must be enabled to view this page.

members
magnitude
magnitudeUnassigned
count
unassigned
taxon
rank
score

psbD.gene2kegg2abundance2tax.norm.forni.trans


7
83.8057193909

superkingdom
4
2
58.203107318
195

58.203107318
195
phylum
4
1117

0

A09100 Metabolism
58.203107318
1
195
3
order
1890424

1890438
2
family
195
58.203107318

47251
2
genus
195

A09100 MetabolismA09100 Metabolism
58.203107318

0
0
subclass
1
1301283

0
0

A09100 Metabolism
1150
1
order

195
25.6026120729
2759
3
superkingdom

0

A09100 MetabolismA09100 Metabolism
25.6026120729
195
2
kingdom
3
33090

35493
phylum
1
195
25.6026120729

25.6026120729

A09100 Metabolism
195
1
subphylum
131221
