## Supplementary Material 1 for "Diel transcriptional pattern contributes to functional and taxonomic diversity in supraglacial microbial communities": psbd.krona.taxa.19_30.html

Javascript must be enabled to view this page.

members
magnitude
magnitudeUnassigned
count
unassigned
taxon
rank
score

psbD.gene2kegg2abundance2tax.norm.forni.trans


7
271.6995

3
2759
195
superkingdom
31.5827

2

A09100 MetabolismA09100 Metabolism
33090
195
31.5827
kingdom
20.2388
3

1
195
35493
phylum
11.3439

131221
195

A09100 Metabolism
subphylum
11.3439
1

superkingdom
240.1168
2
195
4

4
phylum
240.1168
1117
195

1
195
1301283
141.146
subclass

1

A09100 Metabolism
195
1150
141.146
order

13.0792
3
1890424
195
1

A09100 Metabolism
order
98.9708

family
85.8916
1890438
195
2

85.8916
genus

A09100 MetabolismA09100 Metabolism
195
47251
2
