## Supplementary Material 1 for "Diel transcriptional pattern contributes to functional and taxonomic diversity in supraglacial microbial communities": pufM.krona.taxa.4_30.html

Javascript must be enabled to view this page.

members
magnitude
magnitudeUnassigned
count
unassigned
taxon
rank
score

pufM.gene2kegg2abundance2tax.norm.forni.trans


3.19541757965
6

2
2020
superkingdom
3.19541757965
6

2020
1224
3.19541757965
0
6
1
phylum

A09130 Environmental Information Processing

28211
0
class
2
0

204457
0
order
0
1

family
1
0
41297
0

genus

A09130 Environmental Information Processing
0
1
13687
0

0
1
order
0
204441

0
1
family
0
433

50714
0
genus
1
0

1
0
species

A09130 Environmental Information Processing
0
50715

class
3
3.19541757965
28216
2020

80840
2020
order
3.19541757965
3

3.19541757965
3
genus
2020
212743


A09130 Environmental Information ProcessingA09130 Environmental Information ProcessingA09130 Environmental Information Processing
species
3.19541757965
3
1736528
2020
