## Supplementary Material 1 for "Diel transcriptional pattern contributes to functional and taxonomic diversity in supraglacial microbial communities": pufM.krona.taxa.7_30.html

Javascript must be enabled to view this page.

members
magnitude
magnitudeUnassigned
count
unassigned
taxon
rank
score

pufM.gene2kegg2abundance2tax.norm.forni.trans


32.44098330525
6

2020
32.44098330525
6
superkingdom
2

2020
6
phylum
32.44098330525

A09130 Environmental Information Processing
1
18.7289609043
1224

7.58181136605
3
class
2020
28216

80840
2020
order
3
7.58181136605

212743
2020
genus
3
7.58181136605

1736528

A09130 Environmental Information ProcessingA09130 Environmental Information ProcessingA09130 Environmental Information Processing
7.58181136605
species
3
2020

2020
6.1302110349
2
class
28211

204441
0
0
1
order

433
0
family
1
0

0
genus
1
0
50714

50715
0
species
1
0

A09130 Environmental Information Processing

2020
6.1302110349
order
1
204457

41297
2020
1
6.1302110349
family

2020

A09130 Environmental Information Processing
1
genus
6.1302110349
13687
