## Supplementary Material 1 for "Diel transcriptional pattern contributes to functional and taxonomic diversity in supraglacial microbial communities": pufM.krona.taxa.13_30.html

Javascript must be enabled to view this page.

members
magnitude
magnitudeUnassigned
count
unassigned
taxon
rank
score

pufM.gene2kegg2abundance2tax.norm.forni.trans


6
0

0
2
superkingdom
6
0

phylum
0
1224
1
0
6

A09130 Environmental Information Processing
0

0
2
class
28211
0

204441
0
order
1
0

1
433
0
family
0

0
0
50714
genus
1

species
50715
0
1

A09130 Environmental Information Processing
0

0
1
order
204457
0

0
0
41297
family
1

1
genus
0
13687
0

A09130 Environmental Information Processing

28216
0
class
3
0

0
3
0
80840
order

genus
0
212743
3
0

0

A09130 Environmental Information ProcessingA09130 Environmental Information ProcessingA09130 Environmental Information Processing
3
1736528
0
species
