## Supplementary Material 1 for "Diel transcriptional pattern contributes to functional and taxonomic diversity in supraglacial microbial communities": pufM.krona.taxa.19_30.html

Javascript must be enabled to view this page.

members
magnitude
magnitudeUnassigned
count
unassigned
taxon
rank
score

pufM.gene2kegg2abundance2tax.norm.forni.trans


6
115.67751

2020
superkingdom
115.67751
2
6

6
115.67751
1224
2020
1
25.9624

A09130 Environmental Information Processing
phylum

2020
class
82.319
28216
3

3
82.319
80840
2020
order

82.319
212743
3
genus
2020


A09130 Environmental Information ProcessingA09130 Environmental Information ProcessingA09130 Environmental Information Processing
species
2020
3
1736528
82.319

7.39611
28211
2
2020
class

2020
order
204457
1.8104
1

41297
1.8104
1
family
2020

2020

A09130 Environmental Information Processing
genus
1
1.8104
13687

order
2020
204441
5.58571
1

2020
family
1
433
5.58571

50714
5.58571
1
2020
genus

5.58571
50715
1
2020
species

A09130 Environmental Information Processing
