## Supplementary Material 1 for "Diel transcriptional pattern contributes to functional and taxonomic diversity in supraglacial microbial communities": rbcL.krona.taxa.4_30.html

Javascript must be enabled to view this page.

members
magnitude
magnitudeUnassigned
count
unassigned
taxon
rank
score

rbcL.gene2kegg2abundance2tax.norm.forni.trans


46

A09100 MetabolismA09100 Metabolism
2
51.0902807112512
0

4.38422068602

A09100 MetabolismA09100 MetabolismA09100 MetabolismA09100 Metabolism
4
36
superkingdom
51.0902807112512
2
670

1117
0
0
8
phylum
0
4

A09100 MetabolismA09100 MetabolismA09100 MetabolismA09100 Metabolism

subclass
2
1301283
0
0

1150
0
0
order
2

A09100 MetabolismA09100 Metabolism

1890424
0
0
2
order

family
2
0
0
1890431

0
217161
0
genus
2

0
1173032
0
species
2

A09100 MetabolismA09100 Metabolism

22
phylum
2

A09100 MetabolismA09100 Metabolism
0.826243674268
670
1224
46.7060600252312

2008785
670
3.60825315518
class
4

4
order
3.60825315518
119069
670

3.60825315518
206349
670
family
4

4
genus
3.60825315518
70774
670


A09100 MetabolismA09100 MetabolismA09100 MetabolismA09100 Metabolism
species
4
3.60825315518
670
297

0
0
28216
2
class

order
2
0
0
80840


A09100 MetabolismA09100 Metabolism
species
2
0
864051
0

class
14
42.2715631957832
28211
670

356
670
1.6504778913032
order
10
0.4303306276674
4

A09100 MetabolismA09100 MetabolismA09100 MetabolismA09100 Metabolism

family
6
0.436411618554

A09100 MetabolismA09100 Metabolism
2
335928
670
1.2201472636358

4
genus

A09100 MetabolismA09100 MetabolismA09100 MetabolismA09100 Metabolism
670
99
0.7837356450818

4
order
670
204455
40.62108530448

4
family
40.62108530448
31989
670

27.4311099554
2

A09100 MetabolismA09100 Metabolism
4
genus
40.62108530448
265
670

670
147645
13.18997534908
2
species

A09100 MetabolismA09100 Metabolism

2
species

A09100 MetabolismA09100 Metabolism
0
77133
0

8
superkingdom
0
2759
0

0
33090
0
8
kingdom

4
phylum
0

A09100 MetabolismA09100 Metabolism
2
3041
0
0

class
2
0
3166
0

3042
0
0
2
order

family
2

A09100 MetabolismA09100 Metabolism
0
3051
0

phylum
4

A09100 MetabolismA09100 Metabolism
2
0
0
35493
0

2
subphylum

A09100 MetabolismA09100 Metabolism
131221
0
0
