## Supplementary Material 1 for "Diel transcriptional pattern contributes to functional and taxonomic diversity in supraglacial microbial communities": rbcL.krona.taxa.7_30.html

Javascript must be enabled to view this page.

members
magnitude
magnitudeUnassigned
count
unassigned
taxon
rank
score

rbcL.gene2kegg2abundance2tax.norm.forni.trans


A09100 MetabolismA09100 Metabolism
46
193.635449163628
1.67118643096
2

670
2759
17.758147123378
superkingdom
8

670
33090
17.758147123378
kingdom
8

4

A09100 MetabolismA09100 Metabolism
4.840572922058
35493
670
phylum
2
3.2735266643

670
1.567046257758
131221
subphylum

A09100 MetabolismA09100 Metabolism
2

670
12.91757420132
3041
4.08117881578
phylum
2

A09100 MetabolismA09100 Metabolism
4

670
3166
8.83639538554
class
2

2
order
8.83639538554
3042
670

family
670
8.83639538554
3051

A09100 MetabolismA09100 Metabolism
2

670
174.20611560929
2
43.82853133338
4
superkingdom

A09100 MetabolismA09100 MetabolismA09100 MetabolismA09100 Metabolism
36

22

A09100 MetabolismA09100 Metabolism
phylum
2
0
1224
5.77737661873
670

2
class
0
28216
0

order
0
0
80840
2

species
0
0
864051

A09100 MetabolismA09100 Metabolism
2

14
class
670
4.956748924986
28211

10

A09100 MetabolismA09100 MetabolismA09100 MetabolismA09100 Metabolism
order
4
0
356
4.956748924986
670

4.956748924986
335928
670
family
2
1.305230457262
6

A09100 MetabolismA09100 Metabolism

genus
670
3.651518467724
99

A09100 MetabolismA09100 MetabolismA09100 MetabolismA09100 Metabolism
4

order
204455
0
0
4

0
31989
0
family
4


A09100 MetabolismA09100 Metabolism
4
0
0
265
0
genus
2

2

A09100 MetabolismA09100 Metabolism
species
147645
0
0

4
0.820627693744
2008785
670
class

order
0.820627693744
119069
670
4

4
670
0.820627693744
206349
family

670
0.820627693744
70774
genus
4

species
297
0.820627693744
670
4

A09100 MetabolismA09100 MetabolismA09100 MetabolismA09100 Metabolism

1117
114.69536642652
670
4
phylum
53.6224374662
8

A09100 MetabolismA09100 MetabolismA09100 MetabolismA09100 Metabolism

subclass
670
1301283
54.2462355392
2

54.2462355392
1150
670
order
2

A09100 MetabolismA09100 Metabolism

2
670
1890424
6.82669342112
order

2
670
1890431
6.82669342112
family

2
genus
217161
6.82669342112
670

670
1173032
6.82669342112
species

A09100 MetabolismA09100 Metabolism
2

670
9.90484123066
77133
species

A09100 MetabolismA09100 Metabolism
2
