## Supplementary Material 1 for "Diel transcriptional pattern contributes to functional and taxonomic diversity in supraglacial microbial communities": rbcL.krona.taxa.13_30.html

Javascript must be enabled to view this page.

members
magnitude
magnitudeUnassigned
count
unassigned
taxon
rank
score

rbcL.gene2kegg2abundance2tax.norm.forni.trans


0
46
2

A09100 MetabolismA09100 Metabolism
168.37831714082

1.217567748594
115.04792833422
superkingdom
36
4

A09100 MetabolismA09100 MetabolismA09100 MetabolismA09100 Metabolism
670
2

species
2

A09100 MetabolismA09100 Metabolism
77133
0
0

0
0
4

A09100 MetabolismA09100 MetabolismA09100 MetabolismA09100 Metabolism
0
1117
phylum
8

0
subclass
2
1301283
0

0
order
2

A09100 MetabolismA09100 Metabolism
1150
0

0
1890424
0
2
order

0
1890431
0
family
2

0
2
genus
217161
0

0
1173032

A09100 MetabolismA09100 Metabolism
2
species
0

2

A09100 MetabolismA09100 Metabolism
670
1224
phylum
22
113.830360585626
0.667068613508

28211
670
14
class
2.121314860518

2.121314860518
1.099137437354
4

A09100 MetabolismA09100 MetabolismA09100 MetabolismA09100 Metabolism
670
356
order
10

335928
670
2

A09100 MetabolismA09100 Metabolism
6
family
1.022177423164
0.633558505748

0.388618917416
genus
4

A09100 MetabolismA09100 MetabolismA09100 MetabolismA09100 Metabolism
670
99

0
order
4
204455
0

31989
0
family
4
0

0
0
2

A09100 MetabolismA09100 Metabolism
0
265
genus
4

0
0
147645

A09100 MetabolismA09100 Metabolism
2
species

111.0419771116
2008785
670
class
4

670
119069
4
order
111.0419771116

family
4
670
206349
111.0419771116

111.0419771116
70774
670
genus
4

111.0419771116
species
4

A09100 MetabolismA09100 MetabolismA09100 MetabolismA09100 Metabolism
297
670

28216
0
class
2
0

0
80840
order
2
0

2
species
864051
0

A09100 MetabolismA09100 Metabolism
0

53.3303888066
670
2759
superkingdom
8

kingdom
8
670
33090
53.3303888066

0
0
3041
0

A09100 MetabolismA09100 Metabolism
2
4
phylum

2
class
0
3166
0

order
2
3042
0
0

0
3051
0

A09100 MetabolismA09100 Metabolism
2
family

670
35493

A09100 MetabolismA09100 Metabolism
2
4
phylum
53.3303888066
0

53.3303888066
670
131221

A09100 MetabolismA09100 Metabolism
2
subphylum
