## Supplementary Material 1 for "Diel transcriptional pattern contributes to functional and taxonomic diversity in supraglacial microbial communities": rbcL.krona.taxa.19_30.html

Javascript must be enabled to view this page.

members
magnitude
magnitudeUnassigned
count
unassigned
taxon
rank
score

rbcL.gene2kegg2abundance2tax.norm.forni.trans


2
46
0

A09100 MetabolismA09100 Metabolism
166.272889


A09100 MetabolismA09100 MetabolismA09100 MetabolismA09100 Metabolism
superkingdom
4
2
0.962365
670
134.690379
36

15.7788
670
2
species

A09100 MetabolismA09100 Metabolism
77133

86.62984
47.8057
670
8
phylum

A09100 MetabolismA09100 MetabolismA09100 MetabolismA09100 Metabolism
1117
4

35.9426
670
2
subclass
1301283

2
35.9426
670
1150
order

A09100 MetabolismA09100 Metabolism

1890424
order
2
670
2.88154

2
670
2.88154
1890431
family

2.88154
670
2
genus
217161

2
2.88154
670
1173032
species

A09100 MetabolismA09100 Metabolism

31.319374
1.98
670
22
phylum

A09100 MetabolismA09100 Metabolism
1224
2

class
28211
3.170234
670
14

4
356

A09100 MetabolismA09100 MetabolismA09100 MetabolismA09100 Metabolism
order
10
2.324296
670
3.170234

6
0.845938
0
670
335928
2
family

A09100 MetabolismA09100 Metabolism

99
genus

A09100 MetabolismA09100 MetabolismA09100 MetabolismA09100 Metabolism
4
0.845938
670

0
0
4
order
204455

0
0
4
family
31989

265
2
genus

A09100 MetabolismA09100 Metabolism
4
0
0
0

0
0
2
species

A09100 MetabolismA09100 Metabolism
147645

7.27474
670
4
class
2008785

670
7.27474
4
order
119069

family
206349
7.27474
670
4

7.27474
670
4
genus
70774

670
7.27474
4

A09100 MetabolismA09100 MetabolismA09100 MetabolismA09100 Metabolism
species
297

class
28216
670
18.8944
2

80840
order
2
18.8944
670

864051
species

A09100 MetabolismA09100 Metabolism
2
18.8944
670

8
670
31.58251
2759
superkingdom

33090
kingdom
8
670
31.58251

3041
2
phylum

A09100 MetabolismA09100 Metabolism
4
20.84521
13.2795
670

2
7.56571
670
3166
class

2
7.56571
670
3042
order

2
670
7.56571
3051

A09100 MetabolismA09100 Metabolism
family

35493
2
phylum

A09100 MetabolismA09100 Metabolism
4
10.7373
670
10.7373

0
0
2

A09100 MetabolismA09100 Metabolism
subphylum
131221
